# Metabarcoding analysis on European coastal samples reveals new molecular metazoan diversity

**DOI:** 10.1101/253146

**Authors:** David López-Escardó, Jordi Paps, Colomban de Vargas, Ramon Massana, Iñaki Ruiz-Trillo, Javier del Campo

## Abstract

Although animals are among the best studied organisms, we still lack a full description of their diversity, especially for microscopic taxa. This is partly due to the time-consuming and costly nature of surveying animal diversity through morphological and molecular studies of individual taxa. A powerful alternative is the use of high-throughput environmental sequencing, providing molecular data from all organisms sampled. We here address the unknown diversity of animal phyla in marine environments using an extensive dataset designed to assess eukaryotic ribosomal diversity among European coastal locations. A multi-phylum assessment of marine animal diversity that includes water column and sediments, oxic and anoxic environments, and both DNA and RNA templates, revealed a high percentage of novel 18S rRNA sequences in most phyla, suggesting that marine environments have not yet been fully sampled at a molecular level. This novelty is especially high among Platyhelminthes, Acoelomorpha, and Nematoda, which are well studied from a morphological perspective and abundant in benthic environments. We also identified based on molecular data a potentially novel group of widespread tunicates. Moreover, we recovered a high number of reads for Ctenophora and Cnidaria in the smaller fractions suggesting their gametes might play a greater ecological role than previously suspected.

## Introduction

The animal kingdom is one of the best-studied branches of the tree of life ^1^, with more than 1.5 million species described in around 35 different phyla ^2^. Some authors have suggested there may be more than 10 million species of animals, indicating that there is an extensive unknown animal diversity. This hidden diversity may vary according to the animal phyla considered. Not surprisingly, those animal phyla with microscopic representatives (i.e., those animals with a size below 2mm ^3^, also known as micrometazoans ^4^) are suggested to contain most of this potential unknown diversity^3^.

Marine environments cover most of the earth’s surface. More importantly, all metazoan phyla, except onycophorans, have marine representatives, with up to 60% including microscopic members ^5^. Copepods, for instance, are the most abundant multicellular group of organisms on earth ^6^, highlighting the key role of microbial animals in marine ecosystems. Given that the marine benthic meiofauna is also one of the hot spots of alpha-diversity in the biosphere, marine environments thus appear to be ideal sites in which to analyze animal diversity across phyla.

Classical methods to survey animal diversity, such as isolation and morphological identification, might be ineffective to comprehensively analyze micro/mesozooplanktonic ^7^ and meiofaunal diversity ^8^. The microscopic size of the organisms and the wide variety of morphologies makes the identification process tedious and slow, requiring taxonomists with experience in different groups to properly assess the composition of the community and describe new species or groups. Molecular techniques, and especially high-throughput environmental sequencing (HTES), have recently provided a more efficient method to assess and understand ecological patterns in the microbial world ^9^, including metazoans ^8,10–12^. Although, these studies have mainly focused on richness patterns in marine benthic communities or in zooplanktonic communities, with special attention on copepods ^7,13^. Studies of microbial eukaryotes ^14–16^ and even some animal clades ^17^ suggest that HTES could also be used to detect novel lineages. However, such an approach has yet to be applied across the whole animal kingdom.

To obtain a better understanding of the genetic diversity of the different metazoan phyla, and the potential of HTES to quantify diversity and novelty levels, we analyzed a large dataset of ribosomal small subunit (18S rRNA) V4 region tags from European coastal sampling sites in the context of the BioMarKs project, which was designed to analyze the diversity of unicellular eukaryotes. The BioMarKs dataset is based on 137 RNA and DNA samples from six locations ^14,18^ (Fig. S1; Table S1). The use of RNA in this dataset allows analysis that goes beyond the detection of cells or DNA material in the environment, as it provides a window on biological activity. For each sampling site, there is data from both pelagic and benthic environments, with the pelagic samples being divided into different depths and size fractions (Table S2). The large quantity of data, together with the use of a phylogenetically curated taxonomic assignment has provided a global view of genetic diversity across all metazoan phyla. Our data show that 18S rRNA HTES approaches can be used to infer diversity and novelty. Furthermore, we provide evidence that many unsampled lineages remain among animals, and that there are even some potential novel groups. Consequently, greater efforts should be made to sample specific animal groups, especially in benthic environments.

## Results

### Metazoan18S rRNA reference database

An important point to consider when analyzing diversity by metabarcoding is how the taxonomic assignment is done. It is known that the use of GenBank or SILVA as reference databases to perform the taxonomic assignment ^7,8,12,13,19,20^ can be problematic ^21^. The reason is that those databases contain numerous missannotations that affect the final taxonomic assignment. To avoid this problem and to have the best possible taxonomic assignment, we manually constructed a novel phylogenetically curated metazoan 18S rRNA reference dataset.

Our database included 19,364 18S rRNA sequences retrieved from GenBank. The database was curated in a phylogenetic-wise manner, so that each animal phylum had the widest possible representation of internal groups and that each sequence had a clear taxonomic assignment. The resulting database was subsequently used to assign a taxonomic identity to the approximately 1.5 million reads analyzed, providing a holistic and phylogenetically accurate view of the metazoan diversity.

### General abundance and richness patterns of microbial animals

We first analyzed the relative abundance of metazoan reads within the whole eukaryotic dataset. We found that metazoans reads were quite abundant compared to other eukaryotic groups in both the DNA and RNA samples (Fig. 1; Fig. S2). This high percentage of metazoan reads was especially notable in anoxic pelagic environments and in oxic sediments (Fig.1B). Interestingly, metazoan reads were not only abundant in the micro/mesoplankton fraction (68% DNA, 49% RNA of the total eukaryotic reads), but also in the smaller fractions (i.e., the pico/nano fractions which are less than 20um). The presence of a high percentage of metazoan reads in the smaller fractions is especially relevant in the anoxic environment, with 75% of the DNA reads (and 33% of the RNA) being assigned to metazoans.

**Fig. 1:**
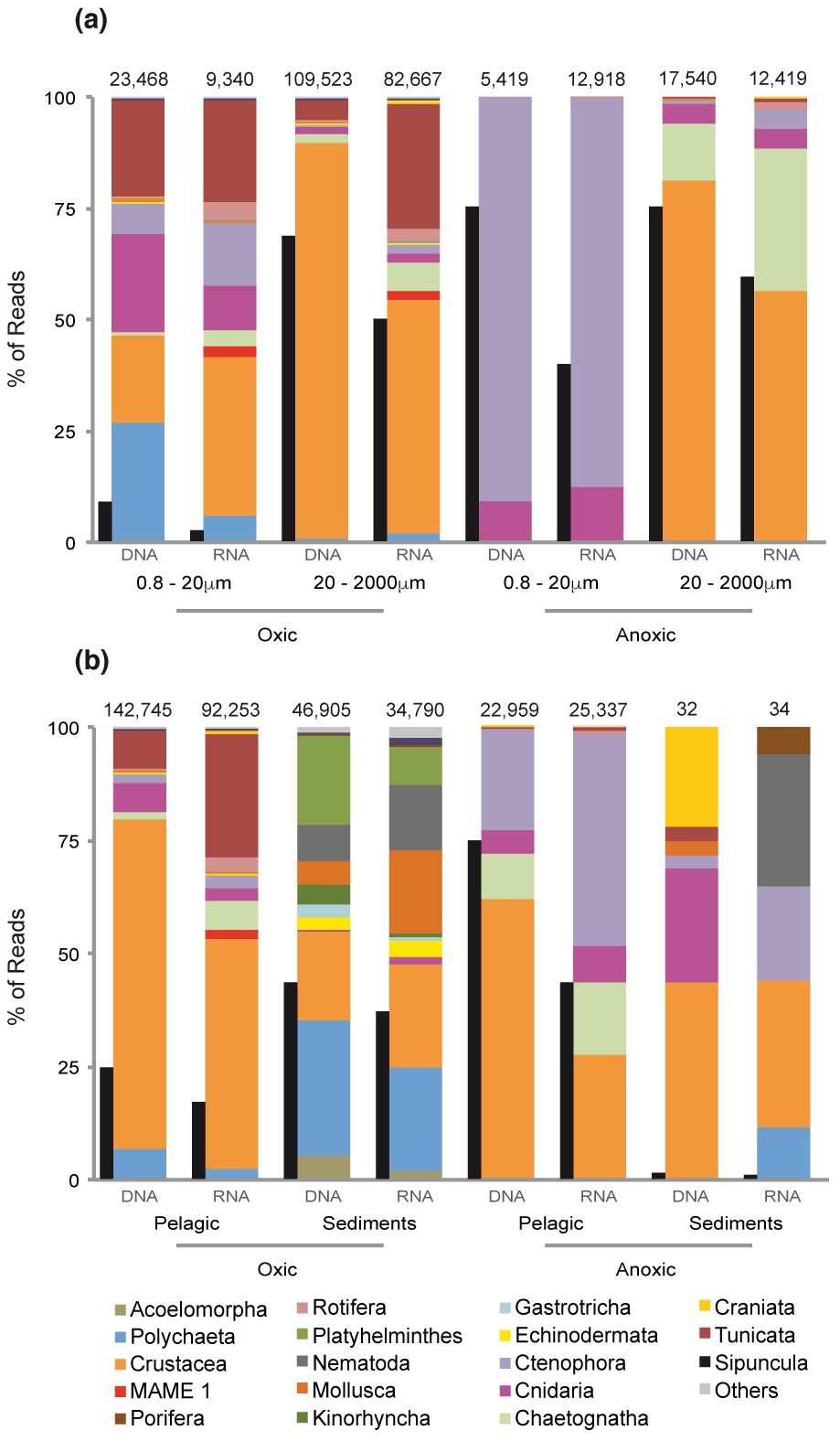
Relative abundances of different metazoan groups and metazoan relative abundance compared to the eukaryotes. Relative abundances of different metazoan groups (colored columns) and metazoan relative abundance compared to total eukaryotes (black columns) in **(a)** oxic fractions and anoxic fractions, and **(b)** different depths, separated by DNA and RNA templates. The number above each column represents the number of metazoan reads in the fraction/environment for the given template (RNA or DNA).

The clustering of reads into OTUs yielded 1067 OTUs from 23 different metazoan phyla (Fig.2, Table S4). 469 OTUs were found to be exclusive to benthic environments, 505 to pelagic environments and 102 OTUs were present in both (Fig.2A). Crustacea appeared as the richest clade (246 OTUs) within the pelagic-exclusive dataset, followed by Polychaeta (45). Within the benthic (sediment)-specific samples, the largest number of OTUs were from Nematoda (227), followed by Crustacea (101). Polychaeta (31) and Crustacea (23) dominated the OTUs present in both environments (Fig.2A).

**Fig. 2:**
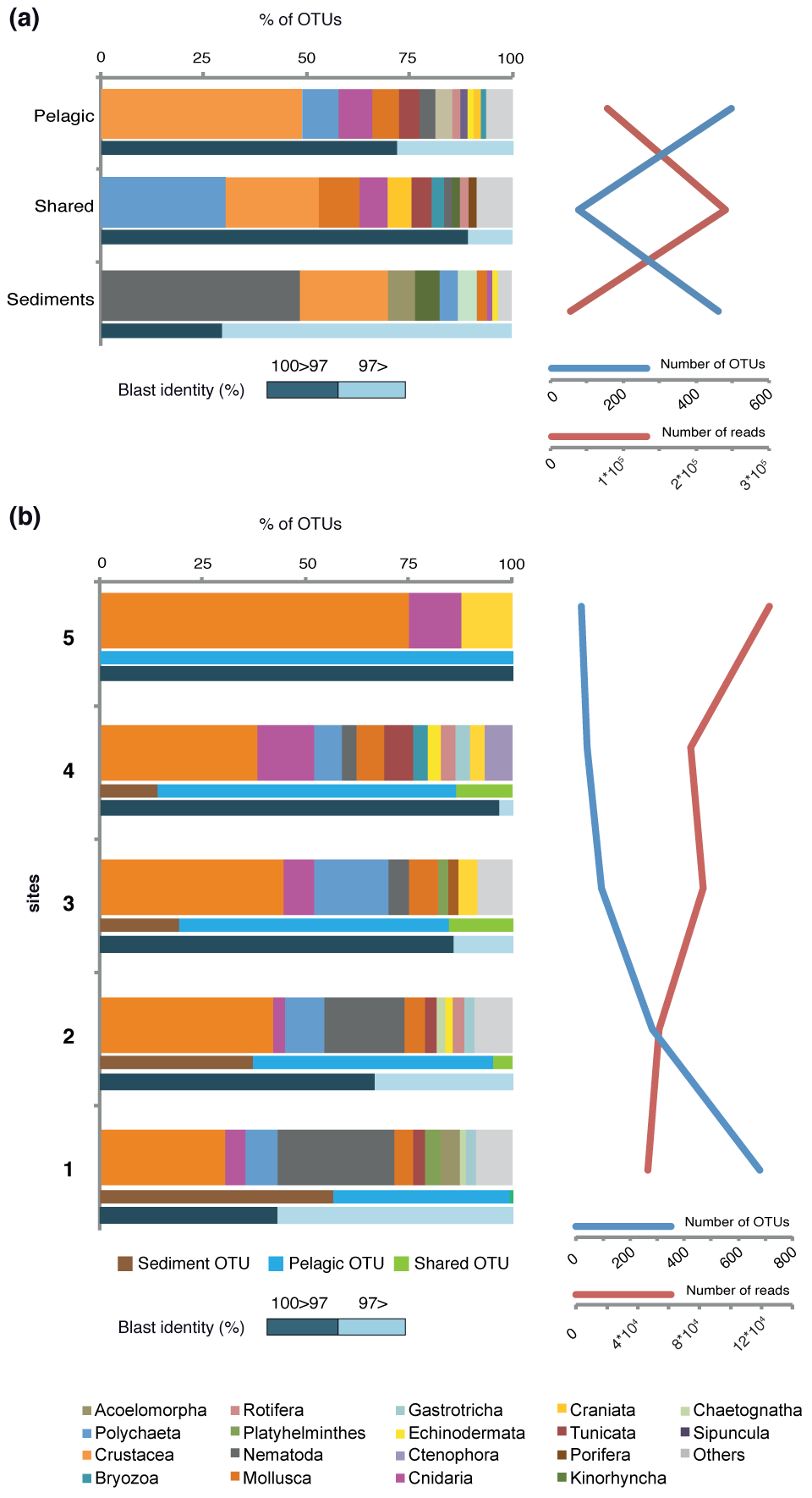
Metazoan richness. **(a)** The OTU distribution for each metazoan group divided into pelagic specific, sediment specific and those present in both environments. BLAST identities are also plotted against NCBI nr nt in dark/light blue. On the right, there is a representation of the number of OTUs (blue line) and number of reads (red line) based on their environment. **(b)** Environmental distribution of OTUs is shown based on prevalence: In blue, pelagic-specific OTUs (i.e., OTU with more than 90% of the reads within the water column); in green, OTUs present both in the water column and the sediments; in brown, OTUs present only in sediments (i.e., OTUs with more than 90% of the reads within the sediments). In addition, BLAST identities are shown against NCBI nr nt in dark/light blue. The number of OTUs (blue line) and number of reads (red line) based on their occurrence in 1 or more (up to 5) geographical site is shown to the right.

The largest proportion of animal reads in oxic water column environments were from Crustacea, which represented up to the 89% of DNA and 53% of RNA in the overall metazoans reads from the micro/meso fractions (Fig. 1A). More than 80% of the crustacean RNA reads, however, corresponded to 8 specific OTUs that were assigned to copepods (Table S5). Besides crustaceans, there was also a high abundance of reads from tunicates (5% DNA only, but 28% RNA) within the oxic micro/mesoplanktonic samples, most of them corresponding to appendicularians (Table S5). On the other hand, benthic samples were dominated by polychaetes (30% DNA, 23% RNA) and crustaceans (19% DNA, 23% RNA) (Fig. 1B). Within benthic Crustacea, ostracods and copepods were the most abundant groups (Table S6).

### Community structure across environments and size fractions

To determine the biogeographical patterns of the microbial animals in our dataset, we analyzed the presence/absence of OTUs in all five sites (discarding the anoxic samples). A large fraction of the OTUs (668 out of 1076) were present in just one single location. However, the number of reads of these "endemic" OTUs (around 4•10^4^) was three times lower than the 8 OTUs present in all sampling sites (around 1.2•10^5^ reads) (Fig. 2B). The taxonomic composition of the cosmopolitan OTUs (Fig. 2B) differed greatly from the complete dataset except for the crustacean dominance (Fig. 2B). In particular, there were no nematodes or polychaetes among the cosmopolitan OTUs, whereas a cnidarian and a craniate OTU appeared to be present over the 5 sampling sites. Our analysis also showed that all the cosmopolitan OTUs belonged to the water column, whereas more than half (56%) of the "endemic" ones belonged to the sediments. These endemic OTUs represented 80% of the total benthic OTUs.

RNA reads indicate metabolically active cells ^22^. Interestingly, we found a relatively high percentage of RNA reads assigned to metazoans in the smaller fractions (from 0.8 to 20 μm): 2.4 % in oxic and 32.4 % in anoxic samples (Fig. 1A). Therefore, we decided to analyze the potential source of those RNA reads. Most of the reads were crustaceans (36% RNA reads), followed by tunicates, ctenophores, cnidarians and polychaetes (Fig. 1A). Ctenophores (85% RNA pico/nano fractions) and cnidarians (16% RNA pico/nano fractions) dominated the reads assigned to metazoans in the anoxic waters of Varna, Black Sea (Fig. 1B).

To understand whether the reads from the smaller fractions were directly derived from the larger ones, we filtered the data based on their co-occurrence between the pico/nano fraction and the micro/meso fractions. We observed that OTUs present in both smaller and larger fractions had a clearly different proportion of reads (Fig. 3). Most of the reads in the smaller fractions belonged to the ctenophores (58%), whereas crustaceans dominated (52%) the micro/mesoplanktonic fractions. In this regard, OTUs corresponding to *Pleurobrachia pileus* (a ctenophore) and *Aurelia aurita* (a cnidarian) were especially enriched in the smaller fraction (Fig. 3), representing 57% of all metazoan RNA reads, and up to 33% of all eukaryotic RNA reads in the anoxic samples (Table S7) (Fig.1A).

**Fig. 3:**
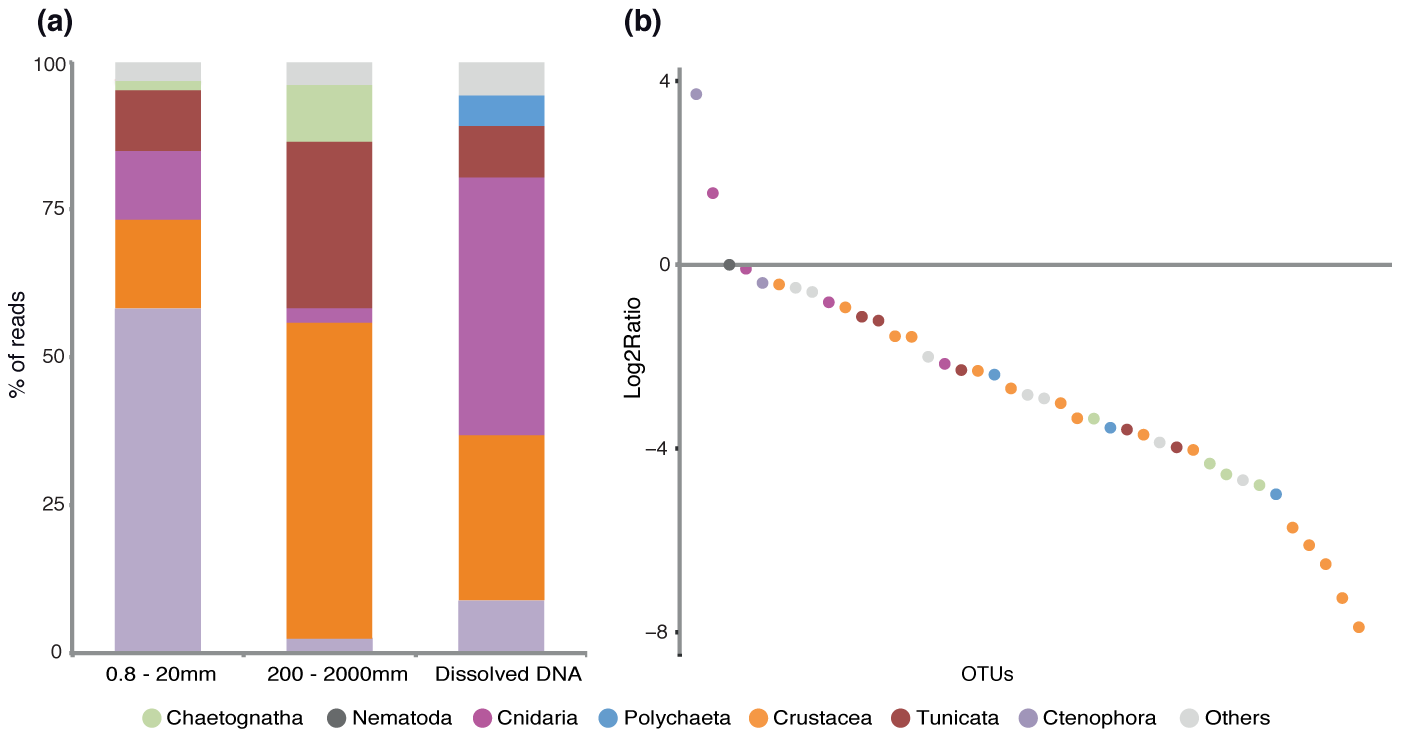
Analysis of the small (pico and nano) and large (micro/meso) fractions, and extracellular DNA. **(a)** Taxonomic distribution of the OTU reads in the smaller and larger fractions and within the extracellular DNA. **(b)** Ratio of the numbers of reads from the smaller fractions and large fraction for these OTUs.

### Sequence novelty

We performed BLAST searches against the NCBI nt nr database to interrogate the level of novelty in our molecular dataset across all animal phyla. The results revealed a high degree of sequence novelty (Fig. 4A). In particular, 35.5% of our OTUs (representing 10.5% of the reads) had a BLAST identity lower than 97% compared to NCBI sequences (Fig. 4B). Moreover, up to 10% of the OTUs, which accounts for 5% of the metazoan reads, had BLAST identities lower than 90%. The putative novelty was especially high among platyhelminthes, acoelomorphs, and nematodes, in which most of their OTUs (75%) had a BLAST identity lower than 97%. Gastrotrichs and crustaceans also had significant novelty (40-50% of their OTUs had a BLAST identity below 97%).

**Fig. 4:**
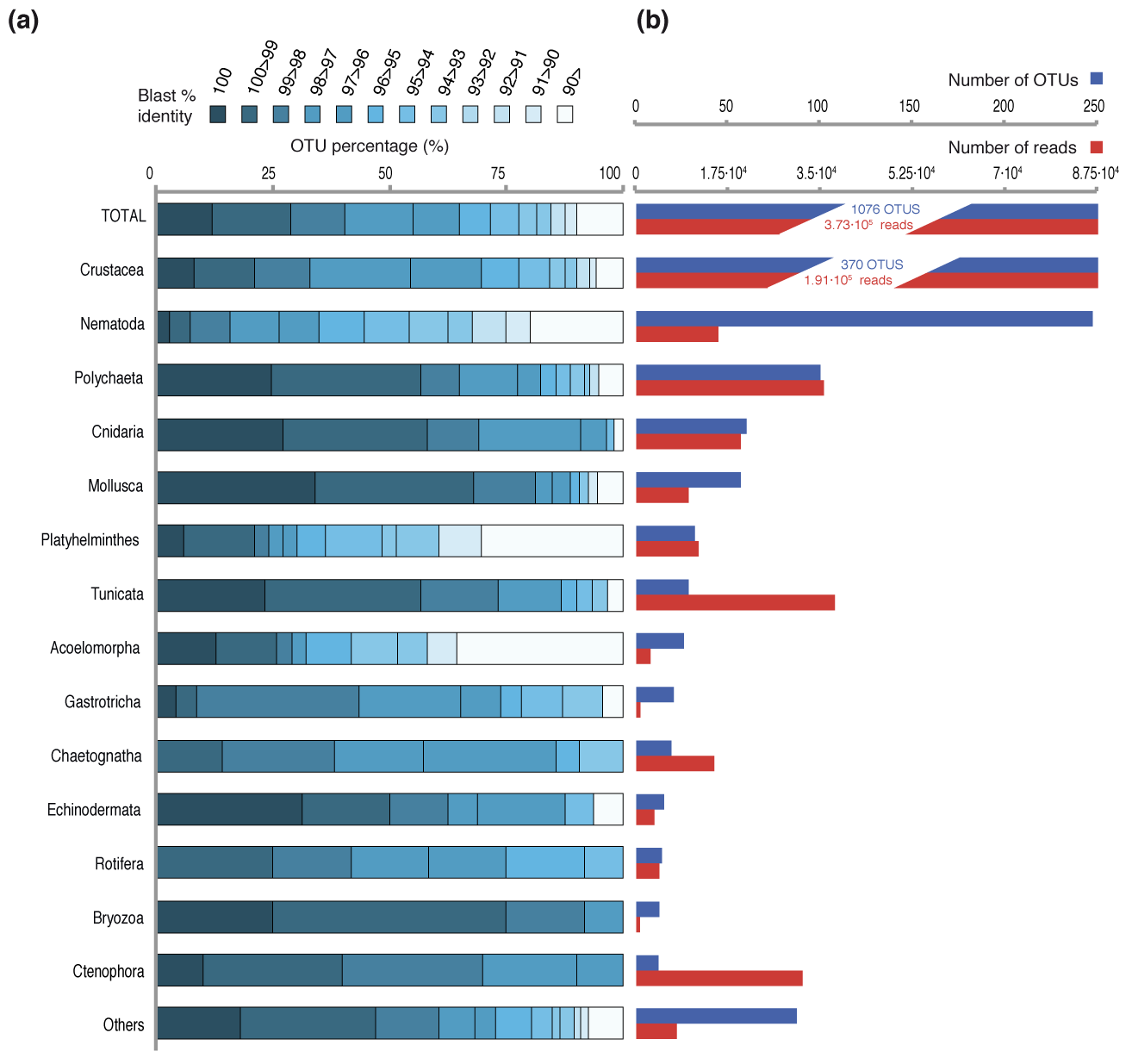
Sequence novelty plus summary of OTUs/read numbers of the main Metazoan phyla in our dataset. **(a)** Distribution of OTU BLAST identities against NCBI nt nr for the main phyla of our dataset. (b) Summary of the number of OTUs (blue) and the number of reads (red) of the given phyla.

Interestingly, the OTUs that appear to be most abundant within the water column (Table S5) and sediments (Table S6) correspond either to already known sequences or with high similarity to known sequences. The level of novelty is also different between benthic and pelagic environments. Thus, 70% of the OTUs found in benthic environments had a BLAST identity of less than 97% (Fig. 2A), while this percentage decreased to 21% of OTUs in the water column or to 11% of OTUs present in both water column and benthos. This suggests that benthic marine environments are a potential hot-spot to find new metazoan taxa or lineages.

Among the potential novelty, we detected a group of three OTUs that had a relatively large number of RNA reads in the water column (1.8%). (Fig. 1, labelled as "MAME 1"; MArine MEtazoan group 1), and with BLAST identities around 95% against two unclassified environmental sequences from GenBank (KC582969 and HQ869055). Analysis of this group of OTUs in other HTES studies based on the 18S rDNA gene revealed 66 more OTUs retrieved from SRA (14 OTUs) and Tara Oceans ^9^ (52 OTUs) that are potentially from the same MAME 1 clade. Those 69 OTUs from BioMarks, SRA and Tara Oceans represent 389,703 reads in total, an indication that OTUs assigned to this group are relatively common in marine environments. Indeed, we found that MAME 1 was present in coastal and open waters with a widespread distribution across the world’s oceans (except for the Arctic) in both the surface and the deep chlorophyll maximum (Fig. S5B).

To have a better understanding of its phylogenetic position, we performed phylogenetic trees. Our trees placed the MAME 1 GenBank sequence within tunicates by both maximum likelihood and Bayesian inference (Table S3), and with good nodal support (79% bootstrap support and 0.99 Bayesian posterior probability), although with relatively longer branches than the rest of the metazoans. To determine its specific phylogenetic position within the tunicates, we inferred an additional tree with most of the available 18S rRNA sequences of tunicates, representing most of the known diversity of this phylum. In this tunicate-focused tree, the MAME 1 sequence clustered with thaliaceans as sister-group to the genus *Doliolium,* although with low nodal support (Fig. 5). Finally, we ran a RAxML-EPA analysis to place the 69 OTUs plus the other NCBI sequence within the reference tree of metazoans and the tree of tunicates. In both cases, the 69 OTUs clustered together, with the reference MAME 1 sequences forming a monophyletic clade. Thus, our phylogenetic analysis suggests that MAME 1 represents a novel, previously undescribed group of tunicates. Given their extremely long-branches, however, additional molecular data will be needed to further confirm this relationship.

**Fig. 5:**
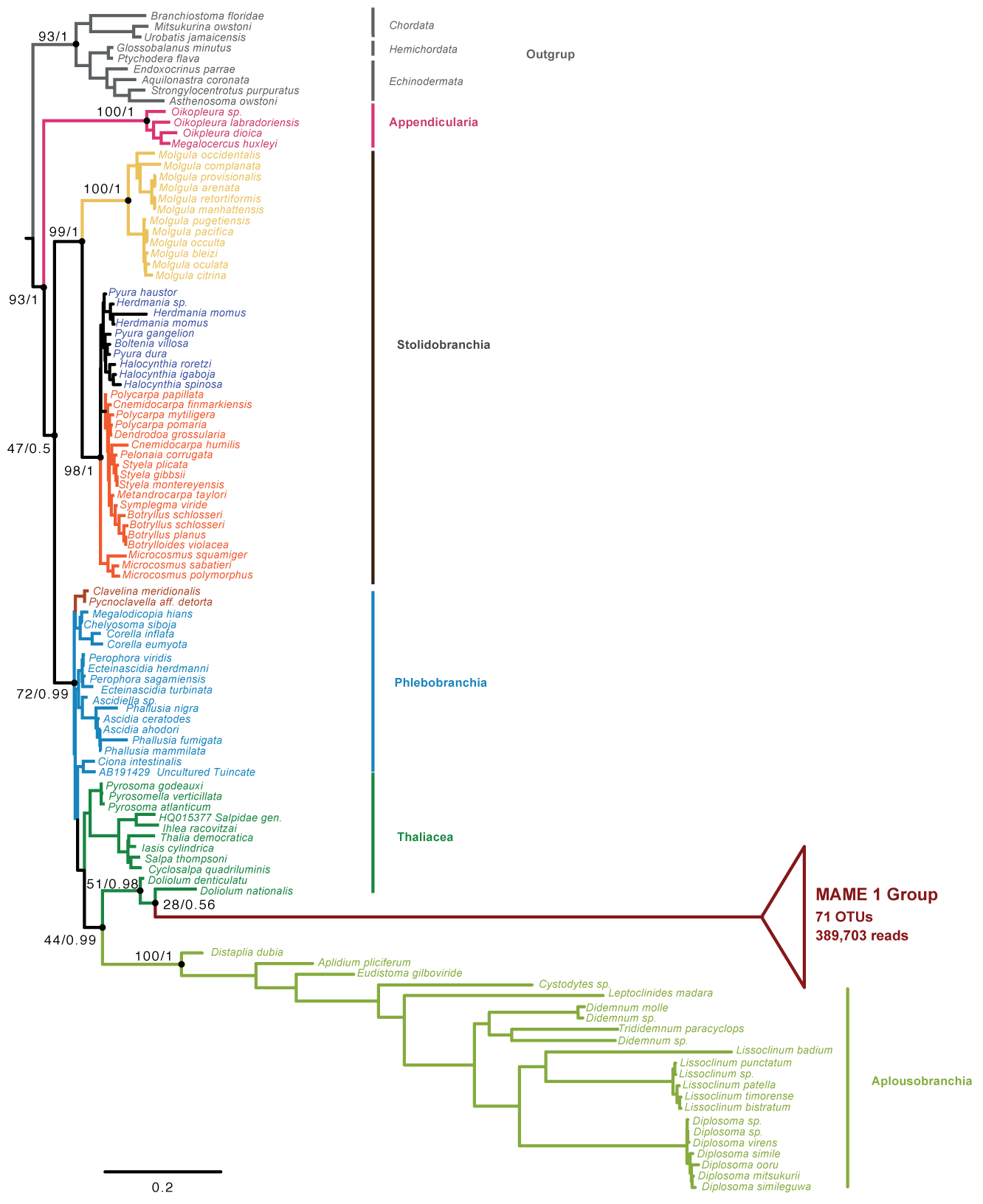
Tunicate 18S rRNA phylogenetic tree placing the novel metazoan group MAME 1. The tree was inferred using RaxML-EPA from the 18S rRNA gene nucleotide sequence and including representatives from all sequenced tunicate groups. The nodal support values marked with a dot correspond to maximum likelihood 100-replicate bootstrap support and Bayesian posterior probabilities.

## Discussion

### High-throughput sequencing, a powerful methodology to assess diversity

HTES is a useful method, but it also has some caveats. For example, it is well known that it may be misleading to directly translate reads and OTU numbers into biomass and number of species, respectively. In particular, the use of amplicon data as a proxy for metazoan biomass abundance has been disputed, also with RNA data ^23^. Different number of rRNA copies in the genomes of different taxa, PCR primer mismatches and amplicon lengths can all affect the correlation between morphological and molecular data ^7,24^. However, some studies have indeed shown positive correlations between read abundances and biomass patterns in bivalve and decapod larvae ^19^ and within copepod groups ^7^. Thus, we believe our approach to biomass abundance, although not perfect, is useful enough to report the most abundant groups. A good indication of our approach is that we recovered the general patterns previously described in micro/mesoplanktonic communities based on morphological observations ^25,26^, in which copepods were found to be predominant within micro/mesoplanktonic communities ^6^ followed by appendicularians ^26^. Moreover, we found a more heterogenic distribution in benthic habitats, which is to be expected considering that sediments are known to harbor most of the metazoan diversity ^5^.

Overall, our data confirms that, although with some caveats, HTES is a powerful tool to assess diversity. In this regard, the construction of a phylogenetically curated database to assign the OTU taxonomy has proven to be crucial for our analysis aimed at describing novelty in different metazoan phyla. Our clustering of OTUs at 97% is likely a conservative approach for metazoans ^27^, and some of our OTUs may indeed represent more than one species. This largely depends on each metazoan lineage and its specific 18S rRNA evolution rate. Moreover, primer bias can affect the detection of some groups, meaning that some taxa can be present in the environment but missing in our dataset ^28^. However, by clustering at 97% we can directly compare the results with the rest of the eukaryotes and get a more stringent output avoiding polymorphisms effects and an overrepresentation of the retrieved diversity.

### Benthic-Pelagic relationship

Analysis of benthic and pelagic metazoan communities in our dataset revealed that most OTUs are exclusively pelagic or benthic, showing few overlaps between the two communities, in agreement with our beta-diversity analyses (Fig. S3, Fig. S4A) and the literature available ^29,30^. Only 10% of OTUs from our dataset were present in both benthic and pelagic communities, and these mainly corresponded to polychaetes, crustaceans, molluscs and cnidarians (Fig. 2A). Among the shared OTUs Polychaeta and Mollusca water column reads probably represent juvenile pelagic stages ^31,32^ while the benthic reads from crustaceans and cnidarians, that are predominantly pelagic, come likely from death organisms or debris.

In addition, our data clearly shows that the pelagic OTUs tend to be present in more sites, while most of the benthic OTUs are restricted to one location. The restricted presence of meiofaunal OTUs has been described previously ^20^. Thus, the distribution in the water column fits more with the consideration that “everything is everywhere” ^33^, probably because pelagic animals have fewer dispersal barriers than do benthic ones ^34^.

### An ecological role for gametes?

Somewhat surprisingly, we observed a high percentage of metazoan reads in the smaller size fractions of most water column samples (Figure 1). This includes, as well, the samples derived from RNA templates, probably indicating a significant biological activity of metazoans in those smaller fractions. We believe it is unlikely that those metazoan RNA reads could come from an extracellular origin because RNA is fragile and quickly degraded by ribonucleases, and its structure is easily affected by both oxygen and water ^35^. Furthermore, the RNA reads from pico/nanoplanktonic fractions contain a different taxonomic distribution compared to the extracellular DNA samples and the micro/mesoplanktonic RNA samples (Fig. 1A and Fig. 3A). Thus, and taking into account the small size reported for certain animal gametes, we hypothesize that a large part of those metazoan reads from the smaller fractions most likely come from metazoan gametes.

This is the case, for example, of the reads from smaller fractions assigned to tunicates, ctenophores, cnidarians and polychaetes, since they all use external fertilization. Ctenophora and Cnidaria, which are not only abundant in DNA reads but also have a relatively high number of RNA reads in the smaller fractions (Fig. 3B), might be a particularly notable example of the importance of gametes in the environment. The co-occurrence of reads in both smaller and larger fractions, the overrepresentation in the smaller ones and the fact that their sperm size is smaller than 5 μm ^36,37^ are good indicators that at least the RNA signal of cnidarians and ctenophores might corresponds to gametes. That will not be the case for the reads assigned to copepods in the smaller fractions. They cannot come from gametes, since copepods use internal fertilization and release eggs larger than 50 μm ^38^. Therefore, the crustacean RNA reads observed in smaller fractions (from 0.8 to 20 μm) are probably the result of cell breakage from larger fractions (Fig. 3A). Finally, we note that some of the OTUs that are exclusively retrieved from smaller fractions could also correspond to sperm from organisms that are larger than 2mm or from benthic fauna with external fertilization and gamete sizes less than 10 μm, such as certain ctenophores and polychaetes (Table S7).

It is worth mentioning that metazoan RNA reads corresponding to germline cells could account, in our data, for as much as 3.2% of the total eukaryotic RNA reads in the smaller fractions (Table S7), and up to 33% of eukaryotic reads in anoxic samples. Thus, their numbers are comparable to those from the unicellular heterotrophic flagellates, which usually reach abundances of up to the 40% of eukaryotic RNA reads in pico and nano plankton ^39^. Thus, and considering those abundances, sperm may play an important ecological role in those environments, particularly in the Black Sea anoxic waters. Further research is needed to assess the effect of sperm in microbial nutrient fluxes, especially during spawning events, when it may represent a passive member of the community eaten by other metazoans or protists from micro-scale fractions.

### Novelty in different metazoan phyla

We performed an analysis on novelty by plotting the pairwise identities of the first BLAST hit against NCBI non-redundant database. This provided a distribution of the "novel" OTUs (those with sequence identities lower than 97% to any NCBI sequence) along different environments (Fig. 2) and for different metazoan phyla (Fig. 4). Interestingly, we found that 45% of our metazoan OTUs had less than 97% identity against the NCBI nt nr database. Why a threshold of 97% for novelty? We believe it is the safest one to detect novelty, although we probably miss a lot of intra-genera or intra-class variation, depending in the animal group. It is worth mentioning, however, that by having a threshold of pair-wise identities below 97%, we avoid any potential intra-individual polymorphic variants ^40^. Therefore, we follow the rationale that OTUs that do not have 100% identities but close (98% or higher) against the first BLAST hit from NCBI non-redundant database, are probably the same taxa (maybe representing intraindividual variations) or very closely related species. In contrast, the OTUs that have a BLAST identity under 97% represent much deeper changes, and so, they clearly represent, at least, different taxa than the ones represented in Genbank. Some OTUs, especially those 10% of our OTUs with pairwise identities against GenBank under 90%, may even represent new clades.

Although one could argue that this degree of novelty might reflect sequencing artifacts, we are confident it is not the case because 1) we have followed a stringent chimera and singletons removal process, 2) the reads are distributed across different samples, and 3) they are not homogeneously distributed among taxonomic groups. In addition, around 80% of our OTUs have RNA reads and their taxonomic distribution is almost identical to the DNA OTUs. So, these novel variants present in the RNA subset are transcribed by active organisms and are less prone to be artifacts or rare variants ^41^.

We are aware that detection of novelty in metazoans just with molecular data is challenging, given that the number of described animal species is larger than the number of 18S rRNA sequences available in public databases (Fig. S7B). Therefore, a novel sequence might belong to a species that has already been described but not yet sequenced. A complete database linking morphological and molecular data is needed to fully solve this issue. However, the 18S rRNA data so far available certainly is a good representation of known animal diversity (Fig. S7B), and we believe our study does indicates which metazoan lineages contain the higher levels of hidden molecular diversity, and so, which are the animal groups needed for a more extensive sampling.

Those animal groups with the higher levels of novelty are not others than crustaceans, nematodes, platyhelminthes, gastrotrichs and acoelomorphs. With the exception of crustaceans, these groups occupy early branching phylogenetic positions within the Ecdysozoa or the Lophotrochoa/Spiralia, or even within the Bilateria ^42^. Moreover, the high genetic diversity in often neglected groups such as Acoelomorpha ^17^ and Gastrotricha ^10^ reveals that these groups need a deeper exploration. We cannot rule out the possibility that the relatively fast evolutionary rates of the 18S sequences from nematodes, acoelomorphs and chaetognaths may have an effect on these low similarity values. In addition, intragenomic variability of the 18S rRNA gene, already described in some metazoan groups such as Platyhelminthes ^43^ or Chaetognaths ^44^, can also contribute to these novelty values. Nevertheless, those are specific, isolated cases. There is certainly extensive genetic novelty in our dataset, suggesting that most acoelomorph, platyhelminth, chaetognath, and nematode species have not yet been sequenced. Some of these hidden animal OTUs occupy key phylogenetic positions, which can help to better reconstruct the metazoan tree of life and unravel the evolution of extant species from the Urmetazoan ^17^.

### A potential novel group of tunicates revealed by HTES

We also recovered and genetically described a potential novel group of tunicates, here named as “MAME 1”. It could be argued that this group represents an already described Thaliacean related to the genus *Doliolum* that happens to have never been sequenced or rare variants of the 18S gene belonging to known species. However, we consider these two options unlikely for several reasons. First, the group seems to be well populated (69 OTUs between our data and public repositories) and present in many environments worldwide, not only in coastal waters (Fig. S5). Moreover, the pairwise identity of the two MAME 1 sequences retrieved from NCBI is about 89%, suggesting is not a single species, but rather an entire group of sequences with high genetic variability, forming an independent clade related to Thaliaceans (Fig. 5). In fact, the nucleotide identity among MAME 1 OTUs is similar as the observed among distant *Aplousobranchia* species (for example, there is an 88% of identity between the 18S rRNA of *Distaplia dubia* and *Diplosoma virens*). Finally, different classes of the 18S rRNA gene have not been reported yet in Tunicates (there are 628 tunicate 18S ribosomal sequences available at Genbank) and the percentage of identity of MAME 1 sequences against described Tunicate species seems too low (78% of identity with the best BLAST hit *Thalia democratica*) for a different 18S rRNA type. In animal groups in which different classes of 18S rRNA gene have been described, such as in chaetognaths, the intra-individual variation among 18S classes lies around 90-93% of identity ^44^. Therefore, we suggest that MAME 1 might corresponds to a new group of tunicates that contains a large number of RNA reads within micro/mesoplankton environments and is present in different habitats. However, without morphological data, we cannot truly discard the possibility that those sequences belong to a molecular divergent group of Thaliacean species, already morphologically described, but without genetic data available. Although this emphasizes the powerful of HTES to assess biodiversity and detect novelty, it also highlights its limitations. Thus, it is crucial to continue and improve the classical screenings of marine diversity, with the aim to link altogether morphological and genetic information in order to better understand the metazoan biodiversity of our oceans.

## Conclusions

We have reported an analysis of micrometazoan diversity in the European coast based on HTES that includes, for the first time, both water column and sediments, oxic and anoxic environments, and both DNA and RNA templates. To assess taxonomy, we constructed a novel reference dataset comprising all animal phyla, which was manually and phylogenetically curated. Our data show general read abundance and richness patterns that partially corroborate previous morphological ^5,6,25,26^ and molecular studies ^8,10,13,19,20,45^. Our data showed a high relative abundance of metazoan RNA reads within pico-nano size fractions (0.8-20 μm), suggesting that the sperm of Ctenophores and Cnidarians plays a relevant ecological role as part of the microbial food network. These results show the potential of HTES techniques as a fast and exhaustive method to approach the study of micrometazoan biomass and diversity patterns.

This kind of data has allowed us to describe novelty values found in different animal phyla. We observed that some animal phyla have much genetic novelty that is yet to be unraveled, including novelty in several well sampled groups such as Crustacea, Platyhelminthes or Nematoda. Our finding of a potential new group of widespread tunicates (MAME 1) highlights the value of phylogenetic approaches to identify novel groups within phyla. The finding of MAME 1 in several HTES datasets could be considered the first step in a reverse taxonomic process ^46^ potentially leading to isolation and detailed description. Overall, our data show that, if we truly want to understand the biodiversity of marine environments, it is important to further sample animal taxa within those environments. To achieve that, we need to have better tools for the genetic screening, and especially for the isolation and morphological characterization of these organisms.

## Materials and Methods

### Sampling, 454 sequencing, curation of the sequences and diversity analysis

During the BioMarKs project (biomarks.eu), samples were collected in six European coastal sites (Fig. S1; Table S1). For sampling collection details, DNA/RNA extraction methods, PCR amplifications, 454 sequencing details and read filtering process see the electronic supplementary material. Processed reads allowed to build a OTU (Operational Taxonomic Unit) table (reads per sample) with usearch v8.1.861 ^47^, using the UPARSE OTU clustering algorithm ^48^, at a threshold of 97% similarity. Afterwards, we used our own metazoan reference dataset (available at figshare https://dx.doi.org/10.6084/m9.figshare.3475007.v1) to assign a taxonomical affiliation to our OTUs. Finally, we removed the putative chimeric metazoan sequences using Mothur‘s Chimera Slayer ^49^ and discarded all the singletons. We determined the degree of novelty of our dataset, by blasting the OTU sequences against NCBI nt nr (September 23 2014). The metazoan OTU table obtained was processed for alpha and beta-diversity analyses using QIIME ^50^. See the electronic supplementary material for details on this section.

### Analysis of the RNA reads from the small fractions

Using QIIME scripts, we binned the OTUs that contain RNA reads within the water column of each sampling site into three different groups: 1) OTUs containing the small fractions (pico/nano), 2) OTUs containing the larger fraction (micro/meso), and 3) OTUs present in both small and large size classes. OTUs representing less than 10 RNA reads per site were discarded.

### Phylogenetic analysis of MAME1 sequence tags

In order to phylogenetically place the short reads assigned to the novel metazoan group (MAME 1) within an animal and tunicate backbone, we performed a RAxML-EPA analysis ^51^ using a metazoan and a tunicate reference tree using the longest putative MAME 1 sequence found by BLAST at NCBI nt nr database (*KC582969*), as a unique MAME 1 representative. Using the MAME1 tree and alignment as a reference we recruited environmental 18S rDNA short reads from SRA and Tara Oceans and used them to perform abundance and distribution analyses (see the electronic supplementary material).

## Data accessibility

Electronic supplementary material that accompanies the online version of this article includes materials and methods and supplementary figures and tables. The complete BioMarks sequencing dataset is available at European Nucleotide Archive (EMBLEBI) https://www.ebi.ac.uk/ena, under project accession number PRJEB9133. OTU tables, 18S metazoan database, MAME 1 group OTU table and phylogenetic trees data (alignments, sequences and trees) are available at Figshare: https://dx.doi.org/10.6084/m9.figshare.3475007.v1.

## Acknowledgment

This work was supported by an Institució Catalana de Recerca i Estudis Avançats contract, two grants (BFU-2011-23434 and BFU2014-57779-P) from the Ministerio de Economia y Competitividad (MINECO), one of which (BFU2014-57779-P) was cofunded by the European Regional Development Fund (FEDER), and a European Research Council Consolidator Grant (ERC-2012-Co -616960) to IR-T. We also acknowledge financial support from the Secretaria d’Universitats i Recerca del Departament d’Economia i Coneixement de la Generalitat de Catalunya (Project 2014 SGR 619). JdC is supported by a Marie Curie International Outgoing Fellowship grant (FP7-PEOPLE-2012-IOF - 331450 CAARL). JP acknowledges support from the European Research Council under the European Union‘s Seventh Framework Program (FP7/2007- 2013) / ERC grant [268513]. The work is part of the EU ERA-Net program BiodivERsA, under the project BioMarKs (Biodiversity of Marine euKaryotes).

### Author Contributions

JdC and IR-T designed and coordinated the study. RM and CdV provided the data. DLE, JdC and JP prepared the 18S metazoan database. DL-E and JdC analyzed the data and prepared the figures. DL-E, JdC, JP and IR-T interpreted the data. DL-E, JdC and IR-T wrote the manuscript, while all authors commented the manuscript.

### Additional Information

Competing financial interests: The authors declare no competing financial interests

